# Genome-wide CRISPR/Cas9 screen identifies MAT2A as a critical host factor for BK Polyomavirus

**DOI:** 10.1101/2023.10.20.563362

**Authors:** Laura Caller, Sophia Ho, Katherine Brown, Liane Dupont, Tjasa Zalatel, Rupert Beale, Paul J. Lehner, Colin M. Crump

**Affiliations:** Department of Pathology, University of Cambridge, Tennis Court Road, Cambridge CB2 1QP, United Kingdom; Cambridge Institute of Therapeutic Immunology & Infectious Disease (CITIID), Jeffrey Cheah Biomedical Centre, Cambridge Biomedical Campus, University of Cambridge, Puddicombe Way, Cambridge, CB2 0AW, United Kingdom; The Francis Crick Institute, 1 Midland Road, London NW1 1AT; UCL Dept of Renal Medicine, Royal Free Hospital, Rowland Hill Street, London, NW3 2PF, UK

**Keywords:** Polyomavirus, CRISPR screen, S-adenosylmethionine, Large T antigen, virus-host interaction, antiviral therapy

## Abstract

BK polyomavirus (BKPyV) is a ubiquitous human pathogen that causes major complications for renal transplant patients (polyomavirus-associated nephropathy) and hematopoietic stem cell transplant patients (haemorrhagic cystitis). There are currently no effective antivirals available against BKPyV. As polyomaviruses are small DNA viruses that express very few proteins and utilise host DNA polymerases for their replication, there is limited possibility of targeting viral proteins for therapeutic intervention. As such, there is increasing interest in targeting host pathways to inhibit these viruses. To identify host genes required for BKPyV infection we have conducted the first genome-wide CRISPR/Cas9 screen in BKPyV infected cells. This led to the identification of methionine adenosyltransferase 2A (MAT2A), which we have validated as a host dependency factor for BKPyV infection. MAT2A is a druggable host enzyme that synthesises S-adenosylmethionine, the principal co-factor and methyl donor within cells. We have found that a small molecule inhibitor of MAT2A is a potent antiviral drug inhibiting BKPyV replication, offering a new therapeutic option for treatment of BKPyV diseases.

## Introduction

First isolated more than 50 years ago, the ubiquitous human pathogen BK Polyomavirus (BKPyV) is a small non-enveloped DNA virus of approximately 45 nm in diameter (Gardner *et al*., 1971; Hurdiss *et al*., 2016). After infection in early childhood BKPyV establishes a life-long persistent infection of the reno-urinary tract, with serology studies suggesting up to 80-90% of the population are infected throughout the course of their lives (Kean *et al*., 2009; Ambalathingal *et al*., 2017). Primary infection is usually asymptomatic or causes a short mild fever, followed by persistent infection and periodic shedding in a patient’s urine throughout their life (Ambalathingal *et al*., 2017). Disease predominantly occurs in immunosuppressed patients, in whom BKPyV reactivation can lead to uncontrolled viral replication and subsequent damage to the reno-urinary tract. Of particular concern are kidney transplant and haematopoietic stem cell transplant patients who can, respectively, often develop polyomavirus-associated nephropathy (PVAN) or haemorrhagic cystitis (HC) (Egli *et al*., 2007; Hingorani, 2016). Therapeutic options to treat BKPyV infection remains limited to general DNA virus antivirals such as cidofovir, which are nephrotoxic (Izzedine, Launay-Vacher and Deray, 2005) and ineffective. Therefore, current treatment for patients suffering from PVAN is reduction of the immunosuppressive drug regimen leading to increased risk of transplant graft loss (Johnston *et al*., 2010). There is therefore an urgent need for a greater understanding of BK polyomavirus-host interactions during infection to identify new therapeutic strategies.

BKPyV entry into cells is mediated by b-series gangliosides such as GT1b and GD1b (Low *et al*., 2006; Neu *et al*., 2013) followed by caveolin-independent endocytosis (Zhao *et al*., 2016). Endosomes-containing virions then appear to be trafficked along microtubules to the endoplasmic reticulum (ER) via a late endosomal recycling pathway (Moriyama and Sorokin, 2008). In the ER, virion disassembly begins with disassociation of VP1 disulphide bonds, reducing virion rigidity and exposing VP2 and VP3 sequences, which in turn hijack the ER associated protein degradation (ERAD) and proteasome pathways (Jiang *et al*., 2009; Bennett, Jiang and Imperiale, 2013). This allows partially disassembled virions to be transported out of the ER to nuclear pores, where the α/β importin pathway facilitates virion import into the nucleus (Bennett *et al*., 2015).

A simple virus of limited coding capacity, the BKPyV genome expresses just 8 viral proteins, none of which have polymerase activity (Abend *et al*., 2009; Nomburg *et al*., 2022). As such, BKPyV is wholly reliant on host DNA replication machinery for its genome replication and production of progeny virions. A genome-wide CRISPR screen offered a convenient and unbiased approach to identify host genes essential for BKPyV infection. Identification of host genes important in BKPyV entry and early gene expression would not only further elucidate the viral entry pathway, but may also determine unidentified, druggable host gene products required for BKPyV infection. Such host genes may be favourable targets for prophylactic drugs in kidney and stem cell transplant recipients, with the aim of preventing reactivation of BKPyV infection.

The use of genome-wide CRISPR screens to increase our understanding of virus infection processes has been highly successful. For example, such screens identified host LDL-receptor (LDLRAD3) as the entry receptor for Venezuelan Equine Encephalitis Virus (VEEV) (Ma *et al*., 2020). In a similar manner TRIM26 was shown to be an essential factor for successful Hepatitis C (HCV) replication in humans (Liang *et al*., 2021). Furthermore, CMTR1 was found to be an essential factor for efficient cap snatching during Influenza A infection (Li *et al*., 2020). Whole-genome CRISPR screens have also been used to investigate differences in host factors required for the entry of wild type and spike deletion mutants of SARS-CoV-2 into host cells (Zhu *et al*., 2021). To date, CRISPR screens have not been applied to the investigation of BKPyV infection, although a genome-wide siRNA screen has been used successfully to investigate BKPyV entry and replication, identifying Rab18 and Syntaxin 18 as important host factors for intracellular viral trafficking (Zhao and Imperiale, 2017).

We have conducted the first genome-wide CRISPR screen for BKPyV replication and have identified several host-dependency factors for BKPyV entry and/or early viral gene expression. Validation of a key hit, methionine adenosyltransferase 2A (MAT2A), by functional gene knockout and chemical inhibition, revealed the importance of this host enzyme for BKPyV infection and replication. MAT2A catalyses the synthesis of S-adenosylmethionine (SAM) from methionine and ATP. SAM is an essential co-factor and methyl donor in a range of host methylation reactions and is required for splicing, gene expression and cellular chromosome replication (Katoh *et al*., 2011). As such MAT2A may be important for a variety of stages of the BKPyV lifecycle. A number of well described small molecule inhibitors of MAT2A activity have recently been developed, some of which are currently in Phase I clinical trials as cancer therapeutics (Li *et al*., 2022), highlighting potential drug-repurposing opportunities for treatment of BKPyV-associated diseases.

## Results

### Generation of Cas9-RPTE/TERT1 cells constitutively expressing active Cas9

To identify host genes important for early stages of BKPyV infection (entry to cells and expression of the early viral gene products), a fluorescence activated cell sorting (FACS)-based genome-wide CRISPR screen was designed and implemented. Human telomerase immortalised renal proximal tubule cells (RPTE/TERT1), a cell type which best mimics the natural site of BKPyV infection, were modified to stably express Cas9 by lentivirus transduction. The established Cas9-RPTE/TERT1 cell line constitutively expresses Cas9, as shown by Western blot (Figure S1Ai). To confirm Cas9 activity, Cas9-RPTE/TERT1 cells were transduced with a lentivirus expressing β2M-specific sgRNA and surface MHC class I levels were quantified by FACS; without β2M MHC class I cannot be transported to the plasma membrane. Greater than 99% of cells demonstrated no detectable MHC class I on the surface of β2M sgRNA transduced cells, confirming efficient Cas9 activity in our Cas9-RPTE/TERT1 cells (Figure S1Aii).

### Genome-wide CRISPR screen identifies host dependency factors for BKPyV infection

After binding and entry, the earliest viral proteins expressed during BKPyV infection are large T antigen (LTAg) and small T Antigen (stAg), of which LTAg protein is easily detected by immunofluorescent staining using commercially available antibodies. Furthermore, lentiviral sgRNA vectors can express fluorescent protein markers such that transduced cells can be readily identified. These two features were exploited in the design of our FACS-based screen using the Bassik Human CRISPR Knockout Library, a lentiviral genome-scale CRISPR library that targets all ∼20,500 protein-coding genes in the human genome with 10 sgRNAs per human gene (Morgens *et al*., 2017). The lentiviral vector also expresses a puromycin resistance gene and an mCherry gene.

Cas9-RPTE/TERT1 cells were transduced with the CRISPR Knockout Library and selected with puromycin for 8 days to remove untransduced cells. Approximately 200 million cells were then infected with BKPyV at a multiplicity of infection (MOI) of 5 infectious units per cell (IU/cell), while at least 50 million were cells left uninfected, to serve as the reference library. Infected cells were fixed, permeabilised and immunostained for the viral early gene LTAg at 3 days post BKPyV infection. These cells were then sorted by FACS to collect mCherry positive cells (representing lentiviral transduced cells) and the bottom 10% of cell population for LTAg signal (representing reduced early viral gene expression) (Figure 1A).

**Figure 1:**
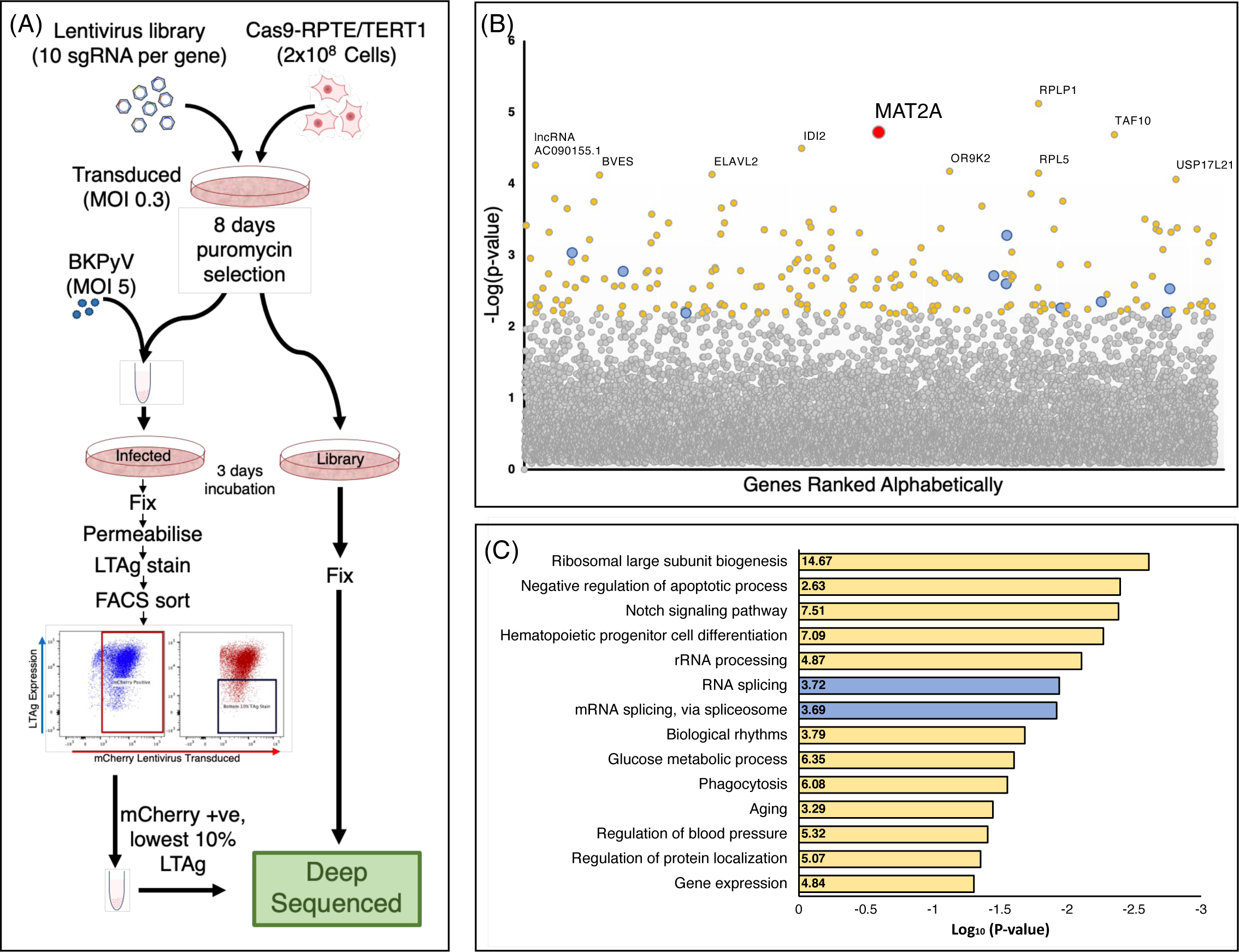
Experimental design of a CRISPR/Cas9 genome-wide screen, identifying MAT2A as a gene important for BKPyV early viral gene expression. (A) *<u>Experimental design of screen</u>*: Cells were transduced with lentivirus comprising sgRNAs of the Bassik CRISPR library, in which there are 10 sgRNAs per gene for all ∼20,500 protein coding genes of the human genome, a puromycin resistance gene and an mCherry gene. Transduced cells were selected in puromycin for 8 days to select for only those cells which were successfully transduced. Puromycin selected cells were then BKPyV infected (MOI 5) or left uninfected to serve as the reference library. After 3 days infected cells were fixed, permeabilised and stained for viral early gene Large T Antigen (LTAg), followed by FACS sorting for mCherry positivity (representing sgRNA transduction and expression), and lowest 10% LTAg expression (reduced early viral gene expression). Next generation sequencing was used to determine the enrichment of specific sgRNA sequences in DNA extracted from the sorted population compared to an unsorted cell population library (B) *<u>Plot illustrating the distribution of genes identified from the CRISPR/Cas9 screen:</u>*All genes quantified are plotted alphabetically (x-axis) against enrichment significance in the lowest 10% of LTAg expressing cells (y-axis), significance displayed as –Log(*p-value*). The top 200 genes (carried forward for DAVID analysis) are either coloured yellow (top 200) or blue (top 200 and involved in RNA or mRNA splicing, a positive control group). Top 10 most significant genes are labelled. One of the most significantly enriched genes was MAT2A, highlighted in this plot in red. Significance of MAT2A *p* < 0.00005, 9 of 10 MAT2A CRISPR sgRNAs were identified. (C) *<u>DAVID analysis of 200 most significantly enriched genes:</u>* The 200 most significantly enriched genes present in the sorted population of cells (lowest 10% of BKPyV infected cells expressing the least LTAg) were subjected to DAVID pathway analysis (v6.8), presented here are the results of functional annotation clustering (EASE score <0.05, Min. count 2).

As the cell population was infected at an MOI of 5 so that >99% of cells should be infected, cells demonstrating lower levels of LTAg are more likely to harbour sgRNA-mediated genetic alterations that disrupt host cell genes required for BKPyV infection. As such, sgRNAs of interest would be enriched in the sorted population when compared to the uninfected and unsorted reference library. In particular, sgRNAs enriched in the sorted population should be specific for genes important in BKPyV entry and early viral gene expression. DNA was extracted from both the sorted “low LTAg” population and the reference library, and sgRNA sequences were amplified by PCR and subjected to Illumina HiSeq. Both library and sorted population reads were trimmed, mapped and quality controlled, showing a balanced high proportion of reads were able to be mapped correctly (Figure S1B).

Screen data were analysed using the MAGeCK method (Li *et al*., 2014), to identify positively selected genes via enrichment of sgRNA sequences associated with these genes in DNA extracted from the sorted population compared with a reference library (Table S1). The enrichment of sgRNA-target genes in the sorted population, compared with a reference library, is displayed with genes plotted alphabetically on the x axis and -Log(p-value) on the y axis (Figure 1B). Most genes fell below a -Log(p-value) of 2 (p>0.01) which we considered as background noise. Approximately 200 genes were significantly overrepresented, scoring above this threshold (Figure 1B; highlighted in yellow, blue, or red). These genes were analysed using the Database for Annotation, Visualisation, and Integrated Discovery (DAVID) to determine the enrichment of any biological processes and pathways (Figure 1C). As expected, RNA and mRNA splicing pathways were identified; LTAg translation can only occur from spliced BKPyV early gene mRNA (Figure 1B and C; highlighted in blue). Another pathway of interest that was identified is Notch signalling, a pathway which regulates cell growth by mediating G1/S progression. Such cell cycle progression is known to be important in BKPyV infection, during which the G1/S check point is bypassed, and a pseudo-G2/M arrest is induced (Caller *et al*., 2019).

Of particular interest was one of the most significant hits identified in the screen: *MAT2A.* This gene encodes for the protein methionine adenosyltransferase 2A (MAT2A). MAT2A catalyses the synthesis of S-adenosylmethionine (SAM) from methionine and ATP. SAM is the methyl donor for a range of host methylation reactions that are involved in the regulation of gene expression and thus MAT2A may be important for various aspects of the life cycle of BKPyV.

### Generation of a MAT2A CRISPR knockout cell line

To validate MAT2A as a host dependency factor for BKPyV infection, we created a stable MAT2A functional knockout cell line by transduction of Cas9-RPTE/TERT1 cells with a lentivirus encoding a MAT2A-specific sgRNA. MAT2A has been shown to be a gene essential for cell survival in haploid human cell models (Blomen *et al*., 2015), thus complete gene function knockout may not be feasible in RPTE cells. Western blot analysis of the puromycin selected population showed clearly reduced, but not abolished, MAT2A expression and so single cell cloning was conducted. Analysis of the selected cell clone, with or without BKPyV infection, demonstrated a significantly reduced signal in Western blot analysis with a MAT2A antibody (which detects two bands), although a faint signal persisted, suggesting a potentially incomplete knockout of MAT2A had been achieved (Figure. 2A). It is important to note that MAT2A also shares a high level of sequence identity with MAT1A (84.3%), and thus many if not all commercially available antibodies to MAT2A are likely to also detect MAT1A, and vice versa. The antibody used for these studies is reported to cross-react with MAT1A. Therefore, it is unclear whether the faint signal observed in our Western blot analyses of MAT2A is due to low level expression of MAT2A or is due to antibody cross-reactivity with MAT1A. Nevertheless, a significant >7-fold reduction in the signal detected with the MAT2A antibody was observed across multiple Western blots, suggesting these cells are not simply lacking one allele of *MAT2A* (Figure. 2B). Immunostaining with a reportedly MAT1A-specific antibody showed no difference in signal intensity for the predominant upper band in Western blots, but reduced signals for the lower band in our MAT2A knockout cells. This suggests MAT1A is expressed in RPTE/TERT1 cells, despite being reported to be a liver specific protein (Torres *et al*., 2000; Ramani and Lu, 2017) and its expression is unaffected by the MAT2A sgRNA sequence used. Furthermore, the MAT1A-specific antibody also appears to cross-react with MAT2A (the lower band) which is greatly reduced in our MAT2A knockout cells. Expression of MAT1A in RPTE/TERT1 cells may help these cells survive in the absence of MAT2A function.

**Figure 2:**
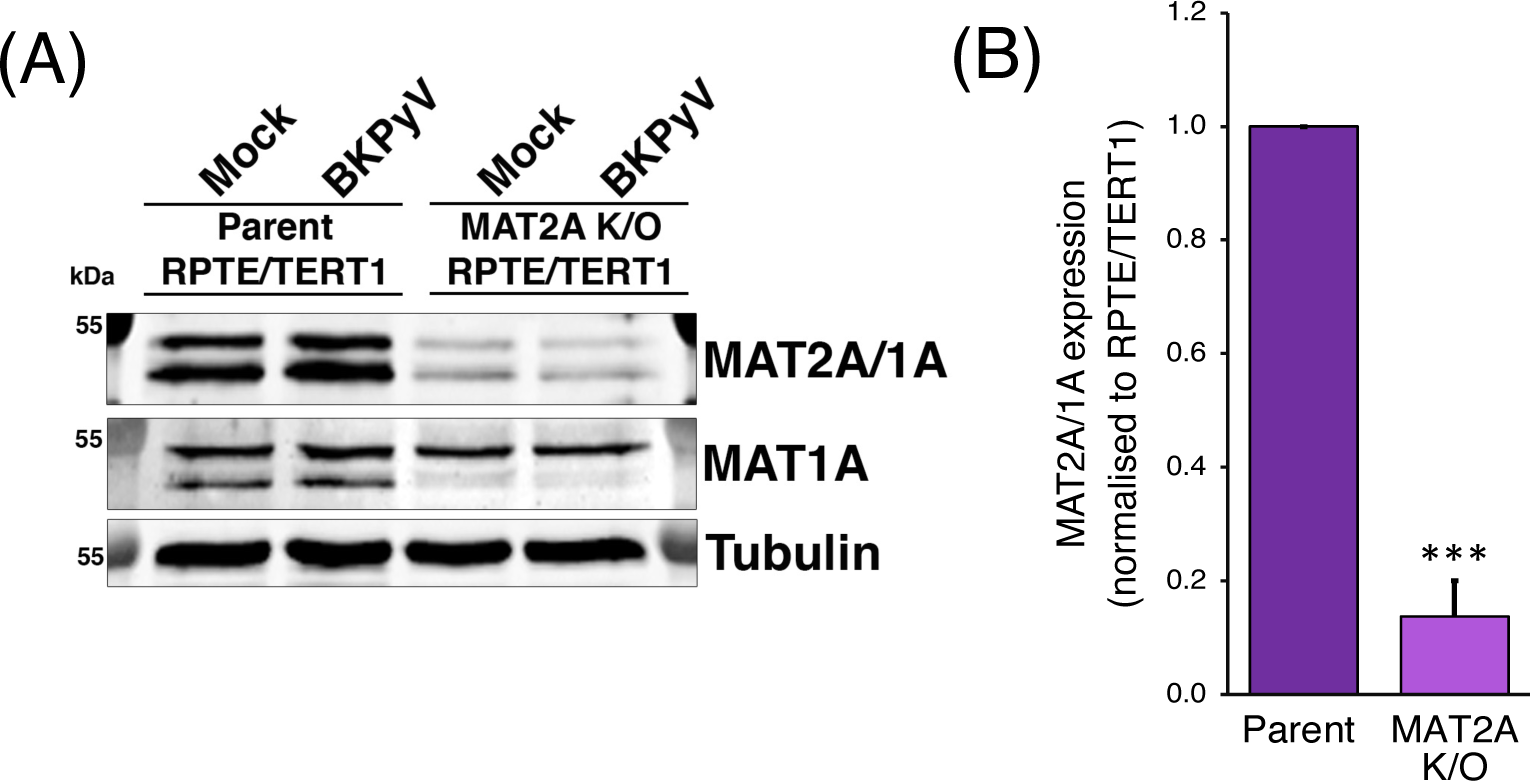
MAT2A Functional knockout RPTE/TERT1 cells show reduced MAT2A expression by Western blot. (A) Parent and MAT2A K/O RPTE/TERT1 cells were infected with BKPyV (MOI 3) or mock infected. At 48 hpi cells were harvested and a Western blot conducted. Stained for MAT2A/MAT1A, MAT1A and tubulin as a loading control. (B) Densitometry of MAT2A/1A (normalized to tubulin) show significant reduction in MAT2A expression in MAT2A K/O cells. Data from three independent experiments, error bars show standard deviation. ** *p* <0.01; students t-test.

Amplification and sequencing of the genomic DNA region from our MAT2A knockout cells encompassing the target site of the sgRNA was used to identify genetic lesions introduced during the CRISPR gene editing process. One MAT2A allele was found to have a single base insertion, introducing a frameshift, and resulting premature stop codon near the N-terminus of MAT2A (after amino acid 56). However, the second allele only contained a single base substitution, causing a change in MAT2A amino acid sequence from alanine at position 55 to glycine (A_55_G; Figure S2A). Alanine to glycine is a relatively modest change to an amino acid side chain (loss of a single methyl group), and so may be predicted to have little impact on MAT2A expression or function. However, this mutation is at a crucial point in the ligand binding pocket, where van der Waals forces act between the hydrophobic alanine at position 55 of MAT2A and the methionine ligand itself (Shafqat et al., 2013). Also, the mutation of a hydrophobic alanine to a neutral glycine may destabilise the protein leading to loss of expression. Western blot data supports this notion, as there is a greater reduction in total MAT2A levels (>7-fold) than would be expected if one allele still expressed full length MAT2A (Figure 2A). It should be noted that mutations at the equivalent alanine residue in MAT1A (e.g. A_55_D) are known to have highly pathogenic effects such as neural demyelination, indicating the importance of an alanine residue at this location (Ubagai *et al*., 1995). Alanine at the equivalent residue is conserved in MAT2A homologs across taxonomic kingdoms from humans to plants to bacteria, further indicating its importance (Figure S2B).

SAM is known to be necessary for a wide range of cellular activities, and MAT2A may be essential for cell viability, as suggested from studies in haploid cell culture models (Blomen *et al*., 2015). Survival of our MAT2A knockout cell line might be due to expression of MAT1A or utilisation of the methionine salvage pathway generating sufficient SAM for essential cellular process. However, we cannot exclude the possibility that our MAT2A knockout cell line expresses a limited amount of MAT2A from the A_55_G point-mutated allele. Nevertheless, considering all the data generated during our studies, our cell line is likely to be a functional MAT2A knockout and any residually expressed of MAT2A has significantly reduced function.

### Knockout of MAT2A function inhibits BKPyV infection

To test the impact of loss of MAT2A activity on early stages of the BKPyV infection cycle we infected the MAT2A functional knockout (MAT2A K/O) cell line and parental RPTE/TERT1 cells with BKPyV (MOI 3) and fixed for immunofluorescence at 48 hours post infection (hpi). Cells were probed for early viral protein expression (LTAg), and the number of infected cells were enumerated as a percentage of total cell numbers (DAPI stain) and normalised to parental RPTE/TERT1 cells (Figure 3Ai). We observed a significant reduction in the number of LTAg-expressing MAT2A K/O cells compared to parental cells. The size of infected nuclei, often seen as an indicator of BKPyV replication and virus assembly, was also significantly reduced (Figure 3Aii). Furthermore, the fluorescence intensity of the LTAg signal in MAT2A K/O cells that were positive was significantly reduced compared to parental cells (Figure 3Aiii, iv).

**Figure 3:**
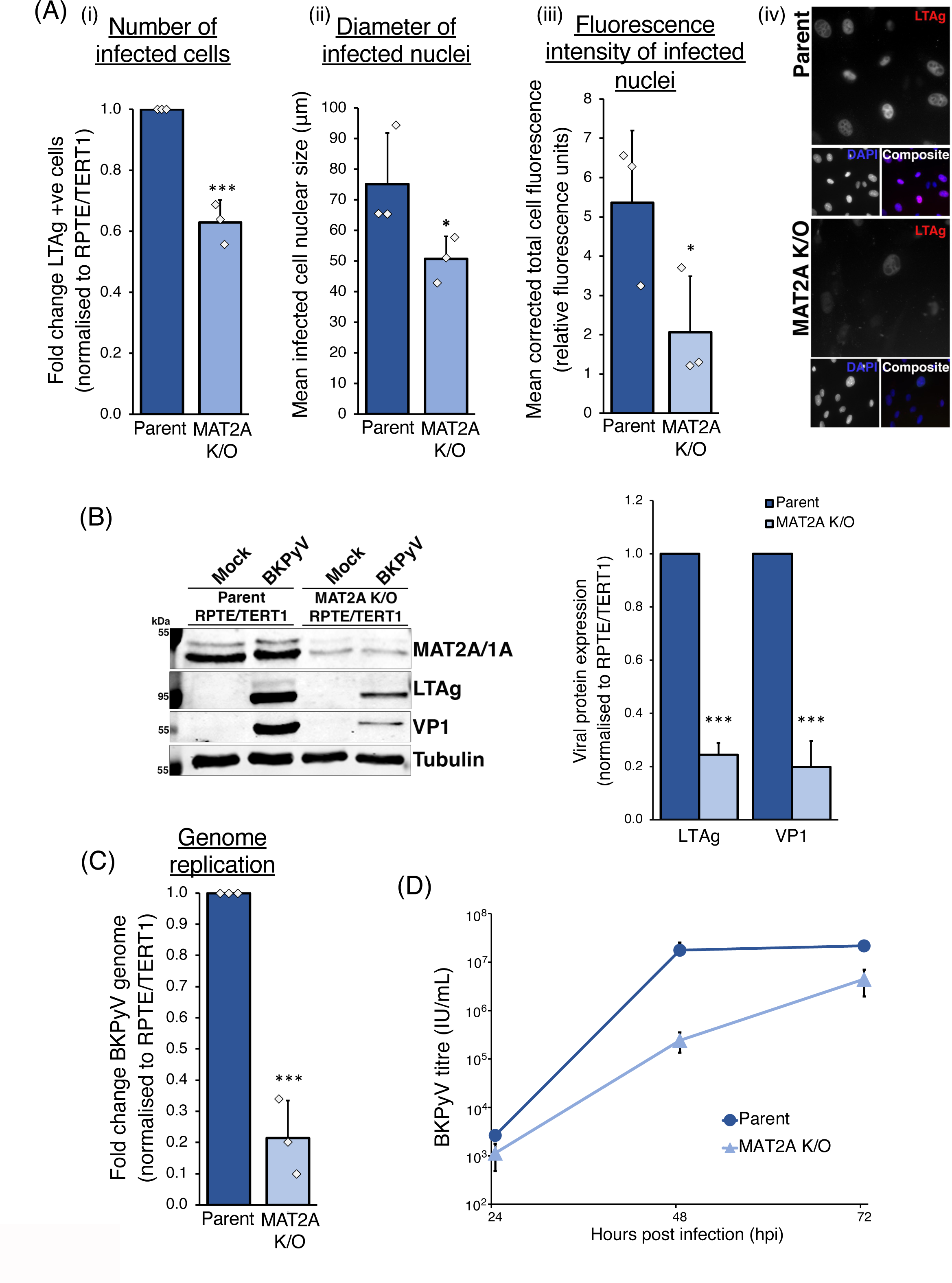
MAT2A Functional knockout RPTE/TERT1 cells validate importance of MAT2A for BKPyV replication. (A) *<u>Effect of MAT2A functional knockout on viral early gene expression</u>*: CRISPR knockout MAT2A single cell clones (RPTE/TERT1 MAT2A K/O) or parent RPTE/TERT1 cells were infected with BKPyV (MOI 3) and formaldehyde fixed at 48 hpi. Cells were stained for LTAg expression and detected by fluorescence microscopy. The number of infected LTAg positive cells were recorded (i), the mean nucleus size of those infected cells was measured (ii) and mean total LTAg fluorescence of infected cells was quantified (iii). All results normalized to parental control cells. Example fluorescence microscopy image shown for reference (iv). (B) *<u>Effect of MAT2A functional knockout on viral early and late gene expression</u>*: Parent and MAT2A K/O RPTE/TERT1 cells were infected with BKPyV (MOI 3) or mock infected. At 48 hpi cells were harvested and a Western blot conducted. Stained for LTAg, viral late gene VP1, cellular MAT2A/MAT1A and tubulin as a loading control. Densitometry of viral proteins LTAg and VP1 (normalised to tubulin) show significant reduction in viral gene expression in MAT2A K/O cells. (C) *<u>Effect of MAT2A functional knockout on viral genome replication</u>*: Parent and MAT2A K/O RPTE/TERT1 cells were infected with BKPyV (MOI 3). At 48 hpi cells were pelleted and qPCR conducted. BKPyV genomes per cell were quantified, using GAPDH as a control. Results normalised to parental control cells. (D) *<u>Effect of MAT2A functional knockout on viral replication</u>:* Parent and MAT2A K/O RPTE/TERT1 cells were infected with BKPyV (MOI 5). At 24, 48 and 72 hpi cells were were harvested and infectious BKPyV was measured by FFU assay. Data from three independent experiments, error bars show standard deviation. * *p* <0.05, ***, *p* <0.001; students t-test.

To investigate the effects on both early and late viral protein expression, MAT2A K/O and parental RPTE/TERT1 cells were either mock or BKPyV infected (MOI 3). At 48 hpi cells were harvested and protein expression levels analysed by Western blot (Figure 3B). A substantial reduction in the levels of both early viral protein LTAg and late viral protein VP1, was observed in MAT2A K/O cells. Quantification of protein levels, normalised to the loading control tubulin, from three independent experiments showed a reduction in LTAg expression of ∼75% and a greater reduction in VP1 expression of ∼80% (Figure 3B).

To investigate the requirement for MAT2A activity for viral genome replication MAT2A K/O and parental RPTE/TERT1 cells were infected with BKPyV (MOI 3), cells were harvested at 48 hpi, DNA extracted and BKPyV genome and cellular housekeeping gene *GAPDH* copy numbers quantified by qPCR. After normalisation to cellular DNA (*GAPDH*) a significant reduction in BKPyV DNA of ∼80% was observed in MAT2A K/O cells compared to parental cells (Figure 3C).

The combined impact of depleting functional MAT2A on viral gene expression in addition to viral genome replication may have a cumulative effect resulting in decreased production of infectious virus. MAT2A K/O and parental RPTE/TERT1 cells were infected with BKPyV (MOI 5) and harvested every 24 hrs up to 72 hpi. Viral titres were determined by fluorescent focus unit assay (Figure 3D). At 48 hpi we observed a >100-fold reduction in the viral titre in MAT2A K/O cells compared to parental cells. The difference in BKPyV titre was reduced to just 8-fold by 72 hpi, likely due to continued slow replication of BKPyV in MAT2A K/O cells, whereas parental cells had reached the maximum level of infectious virus production by 48 hpi.

Taken together this data suggests that loss of MAT2A activity impacts the entire BKPyV lifecycle from early gene expression onwards resulting in significant reductions in virus production.

### Potent inhibition of BKPyV infection by PF-9366, a small molecule inhibitor of MAT2A

MAT2A exists as a homodimer or tetramer, with the methionine and ATP binding active site at each MAT2A dimer interface (Shafqat *et al*., 2013). Additionally, MAT2B is a regulator of MAT2A activity. One copy of MAT2B binds each MAT2A dimer, further activating MAT2A in conditions of low methionine or SAM to induce a rapid increase in the rate of SAM formation.

PF-9366 is an allosteric inhibitor of MAT2A that inhibits binding of the activator MAT2B. A CellTiter Blue assay demonstrated less than 20% reduction on cell metabolism or viability caused by treatment of RPTE/TERT1 cells with PF-9366 at the highest used concentration of 20μM (Figure S3).

To investigate the effects of small molecule inhibition of MAT2A on BKPyV entry and early gene expression, RPTE/TERT1 cells were treated with a range of PF-9366 concentrations, or DMSO control, for 2-3 hours prior to infection. Cells were then infected with BKPyV (MOI 1), and media that contained PF-9366 or DMSO at the appropriate concentrations was replaced at 1 hpi. At 48 hpi, cells were fixed, permeabilised and stained with LTAg-specific antibody and DAPI for detection by fluorescence microscopy. Numbers of LTAg positive cells were counted, correcting for cell numbers (DAPI-positive), and normalised to DMSO control (Figure 4A). A clear dose dependent effect of PF-9366 on the number of LTAg positive cells was observed, which reached statistical significance at concentrations of 5μM and above.

**Figure 4:**
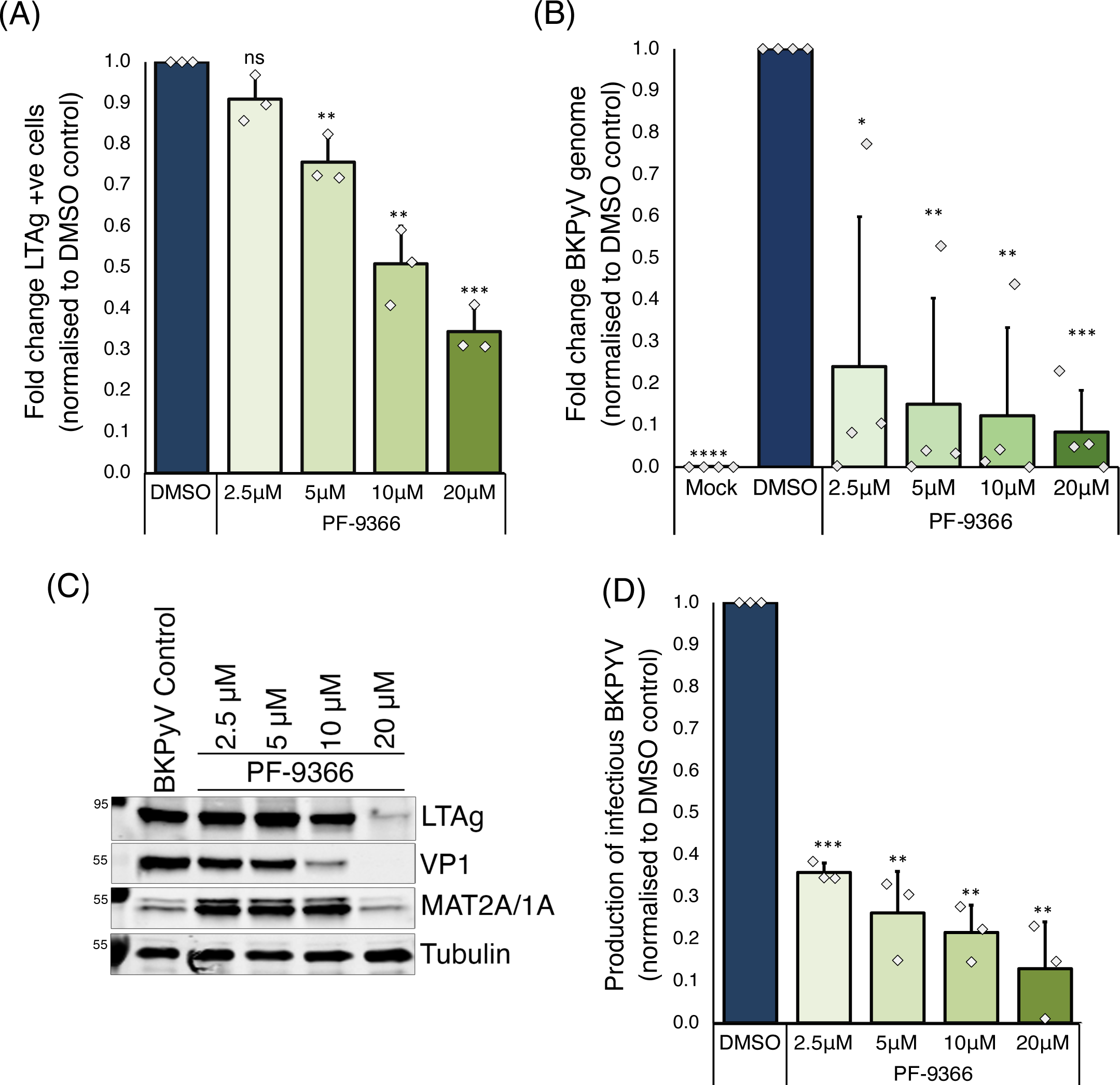
Inhibition of MAT2A with PF-9366 confirms importance of MAT2A activity for BKPyV replication. Parental RPTE/TERT1 cells were treated with PF-9366 at a range of concentrations, or DMSO treated as a control, for 2-3 hours prior to infection. (A) *<u>Effect of PF-9366 on viral early gene expression</u>*: Cells were infected with BKPyV (MOI 1) in media containing appropriate drug concentrations. At 1 hpi infectious media was removed, cells were washed and fresh media with appropriate drug concentrations was added. Cells were formaldehyde fixed at 48 hpi and stained for DAPI and LTAg expression and detected by fluorescence microscopy. The number of infected LTAg positive cells were recorded, normalized to cell number. Data from three independent experiments. (B) *<u>Effect of PF-9366 on viral genome replication</u>*: Cells were infected with BKPyV (MOI 1) in media containing appropriate drug concentrations. At 1 hpi infectious media was removed, cells were washed and fresh media with appropriate drug concentrations was added. BKPyV genomes per cell were quantified, using GAPDH as a control. Results normalised to DMSO treated control cells. Data from four independent experiments. (C) *<u>Effect of PF-9366 on viral early and late gene and cellular MAT2A/1A expression</u>*: Cells were infected with BKPyV (MOI 1) in media containing appropriate drug concentrations. At 1 hpi infectious media was removed, cells were washed and fresh media with appropriate drug concentrations was added. Cells were harvested at 48 hpi and a Western blot performed. Membranes were blotted for viral late gene VP1, LTAg, cellular MAT2A/MAT1A, and tubulin as a loading control. (D) *<u>Effect of PF-9366 on viral replication</u>:* Cells were infected with BKPyV (MOI 1) in media containing appropriate drug concentrations. At 1 hpi infectious media was removed, cells were washed and fresh media with appropriate drug concentrations was added. At 48 hpi cells were harvested and infectious BKPyV was measured by FFU assay. Data from three independent experiments. Error bars show standard deviation. * *p* <0.05, ** *p* <0.01, \*\*\**p* <0.001, \*\*\*\**p* <0.0001, ns = not significant; students t-test.

To explore whether inhibition of MAT2A has an influence on viral genome replication, RPTE/TERT1 cells were treated with a range of PF-9366 concentrations or DMSO control as above and DNA was extracted at 48 hpi. The amount of BKPyV genome and cellular genome present in each sample was quantified by qPCR using primers specific to BKPyV DNA and the cellular housekeeping gene *GAPDH*. Viral DNA was normalised to cellular DNA levels and drug treated samples normalised to DMSO control. These data demonstrated a potent dose-dependent reduction of BKPyV DNA in cells treated with PF-9366, showing a significant ∼4-fold reduction at 2.5μM PF-9366, increasing to >10-fold reduction in BKPyV genome copy number at 20μM PF-9366 (Figure 4B).

Next, the impact of MAT2A inhibition by PF-9366 on early and late viral protein (LTAg and VP1) and MAT2A expression was analysed. RPTE/TERT1 cells were treated with a dose-range of PF-9366 or DMSO as a control and infected with BKPyV as above. At 48 hpi cells were harvested, lysed and protein expression levels determined by Western blot (Figure 4C). Reduced LTAg expression was only observed at PF-9366 concentrations of 10μM and higher, whereas VP1 expression appeared more sensitive to PF-9366 with undetectable levels of VP1 at the highest dose of PF-9366 (20 µM).

Of interest is the effect of PF-9366 on MAT2A expression levels. Even at the lowest dose of 2.5μM PF-9366, a substantial increase in MAT2A expression compared to the untreated control was observed, consistent with published data (Torres *et al*., 2000; Quinlan *et al*., 2017). MAT2A is involved in a feedback loop which regulates available cellular methionine levels, where MAT2A expression increases in response to low cellular SAM levels. Therefore, the reduction in SAM levels caused by PF-9366 inhibition of MAT2A leads to increased expression of MAT2A. This compensation may reduce the impact of PF-9366 at lower inhibitor concentrations. At the highest dose of PF-9366 used (20μM) MAT2A protein levels were similar to DMSO treated control. This suggests that high doses of PF-9366 may prevent de novo synthesis of MAT2A, possibly due to a general suppression of transcription and/or translation.

These data suggest that inhibition of MAT2A has a greater effect on BKPyV genome replication and late gene expression than entry or early gene expression, suggesting that MAT2A activity is important for at additional aspects of the viral lifecycle beyond LTAg expression. The cumulative effect of MAT2A inhibition on the production of infectious virus was assessed by measuring virus titre in the presence of increasing doses of PF-9366. RPTE/TERT1 cells were treated with a dose-range of PF-9366 or DMSO as a control and infected with BKPyV as above. After 48 hours, cells were harvested, and viral titres were assessed by FFU, with data was normalised to DMSO control (Figure 4D). At the lowest dose of PF-9366 (2.5μM) virus replication was significantly reduced by >2.5-fold and viral titres decreased in a dose-dependent manner with ∼10-fold reduction at 20µM PF-9366. The impact of PF-9366 on viral titres was similar to the effect observed on genome copy number (Figure 4B), suggesting the primary effect of MAT2A inhibition by PF-9366 is inhibition of BKPyV genome replication, leading to reduced late gene expression and virion assembly.

### Inhibition of BKPyV replication by PF-9366 is specific to MAT2A function

To confirm the specificity of PF-9366 for MAT2A the impact of PF-9366 on BKPyV replication in our MAT2A functional knockout cells was investigated. MAT2A K/O cells were treated with a dose-range of PF-9366 or DMSO as a control and infected with BKPyV as described above with the exception of using a higher MOI of 3 IU/cell. This is because MAT2A K/O cells were less susceptible to infection than the parental RPTE/TERT1 cells (Figure 3). At 48 hpi cells were either formaldehyde fixed and stained for LTAg and DAPI to investigate virus entry, harvested and DNA extracted for analysis of viral genome replication, or lysed and protein expression levels determined by Western Blot, or viral titres assessed by FFU assays.

No significant change in the number of cells infected with BKPyV, or the expression of LTAg or VP1 was observed in the presence of 2.5µM, 5 µM or 10µM PF-9366 compared to DMSO treated cells (Figure 5A and C). Furthermore, no significant difference in BKPyV genome levels was observed at any concentration of PF-9366 compared to DMSO control (Figure 5B). Similarly, little-to-no difference in the production of infectious BKPyV was observed in the presence of 2.5µM, 5µM or 10µM PF-9366 compared to DMSO treated MAT2A K/O cells (Figure 5D). These data suggest PF-9366 only exerts an inhibitory effect on BKPyV replication when active MAT2A is expressed in cells. Interestingly, MAT2A K/O cells treated with the highest dose (20µM) of PF-9366, did show modest effects (∼2-fold reduction) on the number of LTAg positive cells (Figure 5A) and slightly reduced LTAg protein expression (Figure 5C). Furthermore, more substantial reductions in VP1 expression levels (Figure 5C) and production of infectious titre (Figure 5D) were observed in MAT2A K/O cells treated with 20µM PF-9366.

**Figure 5:**
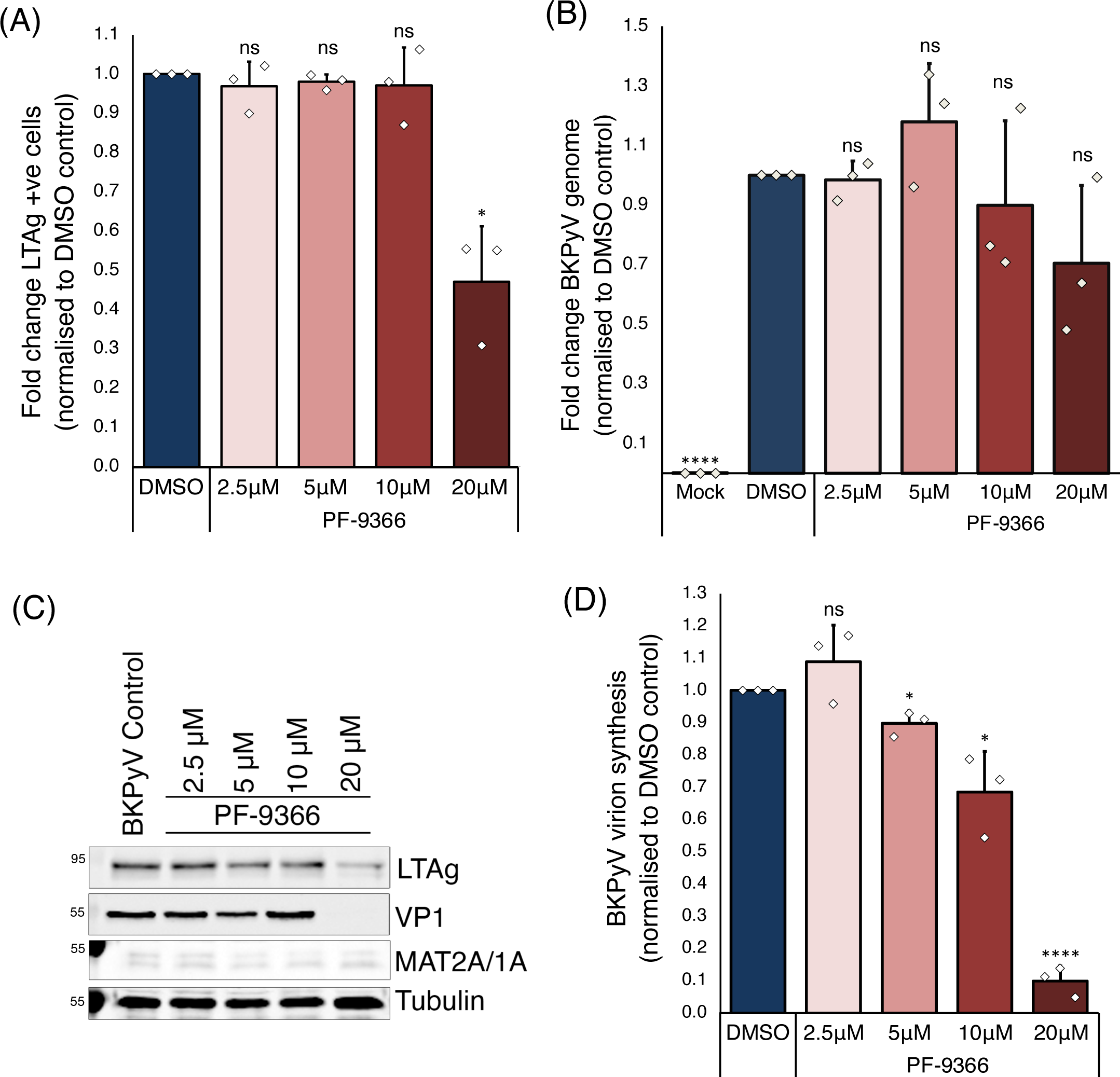
Inhibition of MAT2A with PF-9366 on MAT2A *functional* knockout cells shows minimal effect on virus replication. MAT2A K/O RPTE/TERT1 cells were treated with PF-9366 at a range of concentrations, or DMSO treated as a control, for 2-3 hrs prior to infection. (A) *<u>Effect of PF-9366 on viral early gene expression</u>*: Cells were infected with BKPyV (MOI 3) in media containing appropriate drug concentrations. At 1 hpi infectious media was removed, cells were washed and fresh media with appropriate drug concentrations was added. Cells were formaldehyde fixed at 48 hpi and stained for DAPI and LTAg expression and detected by fluorescence microscopy. The number of infected LTAg positive cells were recorded, normalized to cell number. (B) *<u>Effect of PF-9366 on viral genome replication</u>*: Cells were infected with BKPyV (MOI 3) in media containing appropriate drug concentrations. At 1 hpi infectious media was removed, cells were washed and fresh media with appropriate drug concentrations was added. BKPyV genomes per cell were quantified, using GAPDH as a control. Results normalised to DMSO treated control cells. (C) *<u>Effect of PF-9366 on viral early and late gene and cellular MAT2A/1A expression</u>*: Cells were infected with BKPyV (MOI 1) in media containing appropriate drug concentrations. At 1 hpi infectious media was removed, cells were washed and fresh media with appropriate drug concentrations was added. Cells were harvested at 48 hpi and a Western blot performed. Membranes were blotted for viral late gene VP1, LTAg, cellular MAT2A/MAT1A, and tubulin as a loading control. (D) *<u>Effect of PF-9366 on viral replication</u>:* Cells were infected with BKPyV (MOI 3) in media containing appropriate drug concentrations. At 1 hpi infectious media was removed, cells were washed and fresh media with appropriate drug concentrations was added. At 48 hpi cells were harvested and infectious BKPyV was measured by FFU assay. Data from three independent experiments, error bars show standard deviation. * *p* <0.05, \*\*\*\**p* <0.0001, ns = not significant; students t-test.

Western blot analysis showed no change in MAT2A expression at any concentration of PF-9366 (Figure 5C), suggesting this MAT2A inhibitor is unable to influence the SAM feedback loop that would normally induce MAT2A expression. This is consistent with a lack of MAT2A activity in our MAT2A K/O cell line, suggesting that even if a low level of the MAT2A-A_55_G mutant is expressed in these cells, this mutant enzyme is inactive.

The observed impact of 20µM PF-9366 on VP1 expression and infectious virus production may be due to inhibition of MAT1A, causing reduced viral replication by further reducing cellular SAM levels in our MAT2A K/O cells. PF-9366 has been shown to inhibit MAT1A enzymatic activity by 50% at saturating concentrations. Alternatively, this high dose of PF-9366 may have off-target effects in the MAT2A K/O cells.

Taking all the data from this study together, we have identified MAT2A as an important host dependency factor for BKPyV replication in human kidney epithelial cells. Our data highlights the potential for targeting MAT2A with small molecule inhibitors as a therapeutic avenue to exploit for treating BKPyV disease.

## Discussion

Although the lifecycle of BKPyV has been studied for more than 50 years, we still know little about the host factors and pathways required for successful virus replication. In this study, we have employed the power of genome wide CRISPR/Cas9 screening to identify host pathways involved in the early stages of BKPyV infection, namely up until early viral gene expression. One primary aim of this study was to identify druggable host proteins that are important for BKPyV infection as potential therapeutic targets. Our identification of MAT2A as a host dependency factor for BKPyV replication provides one such target for which small molecule inhibitors have been developed, raising the exciting potential for drug repurposing opportunities to treat BKPyV-associated diseases.

We have validated the requirement for MAT2A activity for BKPyV replication through generating a functional MAT2A knockout cell line and using the small molecule MAT2A inhibitor PF-9366. These orthogonal methods of antagonising MAT2A function both lead to significant reductions in the efficiency of BKPyV infection as well as in early and late viral protein expression, viral genome replication and the production of new infectious virions. This indicates MAT2A activity, and thus presumably methylation reactions, are important at different stages during BKPyV replication. Viral RNA splicing is one process that is likely to be sensitive to MAT2A inhibition and the consequent reduction in SAM levels: the function of splicing regulators can rely on their methylation status, for example SRSF1 methylation by PRMT5 (Radzisheuskaya *et al*., 2019). A complex splicing pattern of both early and late viral mRNA occurs during polyomaviruses infection and is an essential process for the expression of many viral proteins (Nomburg *et al*., 2022). Specific splicing factors were also present within the top hits of our CRISPR screen (e.g., AQR, PTBP2, SFPQ, SRSF11) attesting to the importance of mRNA splicing for BKPyV infection. Translation of viral mRNAs may also be inhibited by reduced SAM levels; methylation of adenosines at the N6 position (m6A) in late viral mRNAs of the related simian polyomavirus SV40 have been shown to enhance viral gene expression (Tsai, Courtney and Cullen, 2018).

Upon sequencing the exon 2 region of the *MAT2A* alleles in our RPTE/TERT1 MAT2A-KO clone to identify the indels near the sgRNA binding site, we discovered that while one allele of *MAT2A* had a frame shift resulting in a premature stop codon that should ablate protein function, the other allele had a single non-synonymous base substitution in codon 55 resulting in a coding change from alanine to glycine (Fig. S2). While this coding change appears to be relatively conservative, our data indicates this is a loss of function mutation. Firstly, MAT2A protein expression in our MAT2A-KO cell line was reduced by greater than 50%, suggesting this A_55_G mutation destabilises the MAT2A protein (Fig. 2). The expression of MAT2A may also be more reduced than is apparent from Western blot data because at least some of the observed signal is likely to arise from cross-reactivity of the antibody used with MAT1A; human MAT1A and MAT2A are both 395aa in length with 84.3% identity and very similar predicted molecular weight (43.6 kDa). Secondly, the MAT2A inhibitor PF-9366 showed no impact on MAT2A expression levels in our MAT2A-KO cell line (Figure 5). Inhibition of MAT2A activity by PF-9366 has been shown to stimulate MAT2A expression (Quinlan *et al*., 2017), likely due to a feedback loop based on cellular SAM concentrations, which we also observed in parental RPTE/TERT1 cells (Fig. 4). This suggests that even if the A_55_G mutant form of MAT2A is expressed, it is not sensitive to PF-9366 and so has presumably lost some enzymatic activity. The side chain of alanine at position 55 in both MAT1A and MAT2A project into the SAM binding pocket and forms van der Waals forces with SAM (Shafqat *et al*., 2013). Furthermore, an alanine at this position is conserved in MAT1A/2A homologues throughout nature, supporting an important role of the methyl side chain of alanine 55 for MAT2A function.

Given the central importance of SAM for all methylation reactions in the cell, it is somewhat surprising that cells with such reduced MAT2A activity are viable. Indeed, this may explain why our MAT2A KO cells may not have a complete knock-out of MAT2A expression and retain a full-length copy of MAT2A with a single amino acid change, which perhaps retains a low level of activity. Alternatively, SAM levels in MAT2A-deficient cells may be maintained by MAT1A, which appears to be expressed in RPTE cells (Fig. 2) despite generally being considered a liver-specific enzyme (Torres *et al*., 2000). Expression of MAT1A in RPTE cells is also consistent with the fact that PF-9366 only demonstrated an impact on BKPyV infection at the highest dose used (20µM); MAT1A is only partially inhibited by PF-9366 (Quinlan *et al*., 2017).

When comparing our CRISPR/Cas9 screen data to that of an siRNA screen on BKPyV infection (Zhao and Imperiale, 2017), we observed surprisingly little overlap in the top hits, with only 3 genes in common within the top 200 hits from both screens (MAT2A, ABCC3, TIMM8B). However, the two screens were conducted using different experimental set-ups, and so it is difficult to draw comparisons on the relative value of CRISPR/Cas9 KO screens versus siRNA knock-down screens without conducting them in parallel with the same experimental conditions and readouts. Despite there being little overlap between the two screens, it is notable that MAT2A is within the top 100 hits from both screens.

In summary, we have conducted the first genome-wide CRISPR screen for host factors required for infection by a human polyomavirus. This has identified MAT2A as an important host enzyme for BKPyV replication. This provides a potential for therapeutic development based on repurposing small molecule inhibitors of MAT2A that have been developed for treating specific (MTAP-deletion) cancers. It will be interesting to investigate whether patients participating in ongoing clinical trials for MAT2A inhibitors show any changes in natural shedding of BKPyV in urine. Our screen data also highlights other host pathways that may be important for BKPyV infection that warrant further investigation. It will be important to conduct additional genome-wide screens for host factors involved in late stages of virus replication to increase our understanding of virus-host interactions and to broaden the range of potential therapeutic targets for treating BKPyV-associated diseases.

## Acknowledgements

We thank V Connor for excellent technical assistance throughout this project, the NIHR Cambridge BRC Cell Phenotyping Hub and their staff for flow cytometry and sorting, and the CRUK Genomics Core Facility for Illumina HiSeq analysis. This work was supported by funding from Kidney Research UK (RP_015_20190306), the Medical Research Council (MR/T016493/1), and the Evelyn Trust (18/48) to RB and CMC. KB was funded by the Wellcome Trust (220814/Z/20/Z, awarded to Andrew Firth). P.J.L. was funded by the Wellcome Trust through a Principal Research Fellowship (210688/Z/18/Z) and the MRC (MR/V011561/1).

## Competing Interests

Some of the work detailed in this manuscript has contributed to a patent filing (Patent number: PCT/GB2023/050875).

## Materials & Methods

### Cell types, virus and primary antibodies

RPTE/TERT1 cells (ATCC CRL-4031) were cultured in DMEM/Hams F-12 supplemented with HEPES (10 mM), hEGF (10 ng/ml), Triiodo-L-thyronine (5 pM), L-ascorbic acid (3.5 μg/ml), holo-transferrin (5 μg/ml), prostaglandin E1 (25 ng/ml), hydrocortisone (25 ng/ml), sodium selenite (8.65 ng/ml), insulin (5 μg/ml), G418 (100 μg/ml), 10 μg/ml gentamicin and 0.25 μg/ml amphotericin B and foetal calf serum (FCS) (0.5%). MAT2A K/O RPTE/TERT1 were grown in the same media except for increased FCS (10%).

BKPyV (Dunlop strain) inserted into pGEM7Zf(+) vector (provided by M. Imperiale, University of Michigan) was prepared, grown, and purified as previously described (Caller et al., 2019).

The primary antibodies used were APC conjugated MHC-I (BioLegend Cat# 311410, RRID:AB_314879), mouse anti-Cas9 (Santa Cruz Cat# sc-517386, RRID:AB_2800509) rabbit anti-SV40 VP1 (Abcam Cat# ab16879, RRID:AB_302561), mouse anti-SV40 T-antigen (Abcam Cat# ab16879, RRID:AB_302561), mouse anti-MAT2A/1A (Santa Cruz Biotechnology Cat# sc-166452, RRID:AB_2266199), rabbit anti-MAT1A (ABclonal Cat# A6650, RRID:AB_2767238) and rat anti-tubulin [YL1/2] (Abcam Cat# ab6160, RRID:AB_305328).

### Cas9 expressing RPTE/TERT1 cell line

To generate Cas9-RPTE/TERT1 cells, RPTE/TERT1 cells at passage 5 were transduced, by spinocculation (250 × g for 1 hour at ambient temperature), with lentivirus generated from the lentiCas9-Blast plasmid, a gift from Feng Zhang (Addgene plasmid #52962; RRID:Addgene_52962) (Sanjana, Shalem and Zhang, 2014), using 16 μg/mL polybrene (Santa Cruz). Three days post transduction, cells were selected with 10 μg/mL blasticidin (Invivogen), refreshed every 3 – 4 days, until all cells in a non-transduced control well had died. Cells were cultured in 5 μg/mL Blasticidin thereafter.

To confirm Cas9 activity, the selected Cas9-RPTE/TERT1 cell population was transduced with a lentivirus generated from pKLV-U6gRNA(BbsI)-PGKpuro2ABFP, a gift from Kosuke Yusa (Addgene plasmid # 50946; RRID:Addgene_50946) (Koike-Yusa *et al*., 2014), containing a β2-microglobulin (β2M) specific sgRNA (5’-GGCCGAGATGTCTCGCTCCG), as described above and selected with puromycin. The selected cell population, along with parental RPTE/TERT1 cells were analysed by flow cytometry for MHC-I expression.

A MAT2A specific sgRNA (FWD 5’-caccgCCTGATGCCAAAGTAGCTTG, REV 5’-aaacCAAGCTACTTTGGCATCAGGC) was cloned into lentiGuide-Puro plasmid, a gift from Feng Zhang (Addgene plasmid #52963; RRID:Addgene_52963) (Sanjana, Shalem and Zhang, 2014). The plasmid was packaged into lentiviruses with packaging plasmid pxPAX2, a gift from Didier Trono (Addgene plasmid # 12260; RRID: Addgene_12260) and pCMV-VSV-G, a gift from Bob Weinberg (Addgene plasmid #8454; RRID: Addgene_8454) (PMID: 12649500) using Lipofectamine 2000 (Thermo Fisher) and Opti-MEM to transfect into HEK293T cells, cultured in DMEM 10% FCS (no antibiotics). Media was changed after 24 hrs and supernatant lentivirus was harvested after another 24 hrs. Lentiviruses were then transduced into Cas9-RPTE/TERT1 cells at passage 14 and single cell cloned. Alteration of the *MAT2A* gene sequence near the sgRNA target site (in exon 2) was confirmed by PCR amplification of the genomic region and by TOPO cloning into pCRII-Blunt-TOPO (Cat# K280002, ThermoFisher), per the manufacturer’s instructions (TOPO primers FWD: 5’-GACTCCCAGGTGCAGAGGGGTT, REV: 5’-TGCTCCAAGGCTACCAGCACGT). DNA extracted from multiple clones were Sanger sequencing.

### BKPyV infections

Cells grown for CRISPR screening were infected with BKPyV at an MOI of 5 fluorescence focus units per cell (FFU/cell) diluted in appropriate medium with 0.5% FCS maximum. At 1 hour post infection (hpi) cells and virus were plated into T150 flasks, diluted further with fresh appropriate medium with 0.5% FCS maximum.

Cells growing in 6-well or 24-well dishes were infected with BKPyV at an MOI of 1 or 3 diluted in appropriate medium with 0.5% FCS maximum. At 1 hour post infection (hpi) medium was removed, cells were washed twice with PBS, and fresh appropriate medium with 0.5% FCS maximum was replaced.

### MAT2A Inhibitor treatments

PF-9366 (Sigma Cat #PZ0403) was dissolved in DMSO to a stock concentration of 20 mM. Cells were treated for 2-3 h prior to infection with the stated concentration of PF-9366 diluted in media. Media containing the stated concentrations of PF-9366 was used during virus inoculation and in final growth media added to cells.

### Flow cytometry

Cells were harvested and blocked in PBS + 2% FCS for 30 mins on ice. Cells were then stained with APC-conjugated antibody to MHC-I for 1 hr on ice, washed with PBS + 2% FCS, fixed in ice cold 3% ultrapure formaldehyde in PBS for 20 mins on ice. Cells were washed with PBS + 2% FCS and then PBS and then analysed using a BD LSR Fortessa flow cytometer (Becton Dickinson). Data were processed using FlowJo™ Software.

### FFU assays and immunofluorescence microscopy

Fluorescent focus unit (FFU) assays were used to determine the concentration of infectious virus in experimental samples (titration FFU assay), or to determine the number of infected cells in certain experimental conditions (infectivity FFU assay).

In titration FFU assays RPTE/TERT1 cells were infected with a 10-fold dilution series of the samples to be titrated, fixed at 48 hpi and immunostained for VP1 expression as described in (Caller *et al*., 2019). In infectivity FFU assays parent or MAT2A K/O Cas9-RPTE/TERT1 cells were untreated or treated with PF-9366 for 2-3 h prior to infection. Cells were then infected in the presence or absence of the different PF-9366 concentrations. At 48 hpi cells were fixed and stained for LTAg expression.

For immunofluorescence analysis, cells were fixed in 3% formaldehyde at room temperature for 15 mins. Fixed cells were then permeabilised and quenched at room temperature for 15 mins (50 mM NH_4_Cl and 0.1% Triton X-100 in PBS), blocked in PGAT at room temperature for 15 mins (0.2% gelatin, 0.01% Triton X-100, 0.02% NaN_3_ in PBS), and stained with primary antibodies diluted in PGAT at room temperature for 1 hr. Secondary antibodies used for immunofluorescence were Alexa Fluor 568-conjugated donkey anti-mouse or goat anti-IgG1 mouse and Alexa Fluor 488-conjugated donkey anti-rabbit or goat anti-IgG2a mouse. Coverslips were mounted using SlowFade Gold with 4′,6′-diamidino-2-phenylindole (DAPI) (Invitrogen). Samples were analysed with an inverted Olympus IX81 widefield microscope. Illumination was performed with a Lumen 200 arc lamp (Prior Scientific) and bandpass filters for DAPI (excitation of 350/50 nm and emission of 455/50 nm), Alexa Fluor 488 (excitation of 490/20 nm and emission of 525/36 nm), and Alexa Fluor 568 (excitation of 572/35 nm and emission of 605/52 nm) (Chroma Technology Corp). Images were acquired with ImagePro Plus software (Media Cybernetics), a Retiga EXi Fast1394 interline CCD camera (QImaging), and a 60 Plan Apochromat N oil objective (numerical aperture 1.42) (Olympus) for a pixel resolution of 107.5 nm/pixel.

To determine numbers of LTAg positive cells in each experiment 10 fields of view were captured for each experiment (5 sites per well, 2 wells per experiment), positive LTAg nuclei were determined by averaging individual image counts and multiplying by 341.9, which corresponds to the number of fields of view per well in a 24-well plate.

When conducting LTAg fluorescence intensity and nucleus size analysis, 20 fields of view were captured for each experiment (5 sites per well, 4 wells) to give a total of n = 40 fields of view per condition, per experiment. All quantification analyses were carried out using ImageJ and an optimised robust automated detection pipeline (Schindelin *et al*., 2012), which came with a benefit of overcoming the limitations of human error and unconscious bias. For each individual experiment, threshold, brightness, and contrast were adjusted and standardised to produce comparable levels of accuracy to manual counting. On average, 48 – 309 nuclei were visualised and analysed per field-of-view. LTAg staining patterns across each field of view were measured by quantifying the fluorescence intensity using the Mean Grey Value (MGV) function of ImageJ and averaging the results for each experimental condition. Individual cell fluorescence intensity was determined by drawing polygon selections around each LTAg-positive nuclei in ImageJ and measuring the nuclear area, MGV and integrated density, then doing the same for background areas close to each nuclei measured. Corrected cell fluorescence was measured by the following formula:

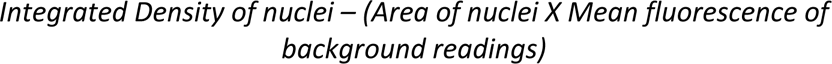

Individual nuclei size was determined by measuring the diameter of each LTAg-positive nuclei in ImageJ.

For ease of interpreting and comparing the effect of MAT2A inhibition or MAT2A functional knockout on BKPyV infection levels and titres, the control condition in each experiment was arbitrarily set to 1, and tested conditions were corrected to the control for each independent experiment. A two-sample *t* test was conducted to test for statistical significance (*P* values) (standard deviations are shown with error bars).

### Western blotting

All cells were lysed by suspension in modified radioimmunoprecipitation assay (mRIPA) buffer (50 mM Tris, pH 7.5, 150 mM NaCl, 1% sodium deoxycholate and 1% Triton X-100) supplemented with Complete protease inhibitors without EDTA (Roche). Cellular debris was removed by centrifugation at 17,000 × *g*. Proteins were separated by SDS-PAGE and transferred to nitrocellulose membranes before being blocked in 5% skimmed milk powder in PBS.

Following primary antibody binding, Li-Cor IRDye 680-conjugated (anti-mouse, anti-rabbit, or anti-rat) or IRDye 800-conjugated (anti-mouse or anti-rabbit) secondary antibodies were used. Membranes were then imaged on a Li-Cor Odyssey infrared imaging system.

Western blot densitometry was conducted Image Studio (LI-COR Biosciences), normalising to tubulin loading control.

### BKPyV DNA quantification by qPCR

To isolate DNA, cells were harvested by scraping into media, pelleted by centrifugation (6,000 × g, 5 mins, 4°C), washed with PBS and further pelleted by centrifugation, then resuspended in 10μl Taq PCR buffer containing 1μg/μl Proteinase K and 2.5 mM MgCl_2_. Cells were heated to 60°C for 1 hr, followed by inactivation of Proteinase K at 95°C for 15 mins. Samples were then diluted to 100μl with PCR grade water.

Amplification and quantification of BKPyV and GAPDH control DNA were performed using a 2× SYBR Green mastermix containing 2.5mM MgCl_2_, 400μM dNTPs, 1/10,000 SYBR Green (Molecular Probes), 1M Betaine (Sigma), 0.05U/μl of Gold Star polymerase (Eurogentec), 1/5 10X Reaction buffer (750 mM Tris-HCl pH 8.8, 200 mM [NH4]_2_SO_4_, 0.1 % [v/v] Tween 20, Without MgCl_2_), and ROX Passive Reference buffer (Eurogentec). Primers were designed and obtained through TIB MolBiol BKPyV (FWD: 5’-TGT CAC GWM ARG CTT CWG TGA AAG TT-3’; REV: 5’-AGA GTC TTT TAC AGC AGG TAA AGC AG-3’), GAPDH (FWD: 5’-GTC TCC TCT GAC TTC AAC AGC G-3’; REV: 5’-ACC ACC CTG TTG CTG TAG CCA A-3’). DNA amplification conditions comprised preincubation at 95°C for 10 min, followed by two steps (40 cycles) at 95°C for 15 s and 65°C for 1 min using VIIA7 Applied Biosystems StepOne Real-Time PCR System.

### CRISPR screen

Cas9-RPTE/Tert1 cells were lentiviral transduced (Sanjana et al., 2014) with a Human CRISPR Knockout library, a gift from Michael Bassik (Addgene #101926 – 101934) at an MOI of ∼0.3 (Morgens et al., 2017). To remove un-transduced cells, starting at ∼36 hr post transduction cells were selected in 10% FCS media supplemented with puromycin (12.5 mg/ml) for 8 days.

At 9 days post transduction, surviving cells were transferred to 0.5% FCS media and either infected with BKPyV (MOI 5) in suspension or left uninfected as a reference library.

At 12 days post transduction (72 hrs post BKPyV infection) cells were harvested into a single cell suspension by trypsinisation (15 min, 37°C). To prevent adherence of cells to one another by DNA leakage the suspension of cells was incubated with 600 Units of DNAse I (Roche) (30 mins, 37°C), with gentle agitation. Cells were the, pelleted by centrifugation (400 × g, 5 mins, 4°C), washed with ice cold PBS, and once again pelleted (400 × g, 5 mins, 4°C).

Cells were then fixed in ice cold 1% paraformaldehyde (PFA), incubated for 20 mins on ice. When fixed cells were pelleted by centrifugation (800 × g, 5 mins, 4°C), washed once again in ice cold PBS, followed by further pelleting.

Cells were then permeabilised with 100% ice cold methanol, incubated for 15 mins on ice. Once permeabilised cells were diluted in 2% foetal calf serum in PBS (FCS/PBS) and centrifuged to pellet cells (800 × g, 10 mins, 4°C), followed by washing with 2% FCS/PBS and further pelleting by centrifugation (800 × g, 10 mins, 4°C).

Cells were then immune-stained at room temperature. Cells were incubated in primary antibody, anti-LTAg PAb416 (Abcam) diluted 1:200 in 2% FCS/PBS, with periodic gentle agitation for at least 1 hr. Cells were then diluted in 2% FCS/PBS and pelleted by centrifugation (800 × g, 10 mins, room temperature), and further washed in 2% FCS/PBS. After once again pelleting by centrifugation cells were then incubated for at least 30 mins with goat anti-mouse 647 secondary antibody (Alexa-Fluor), diluted 1:200 in 2% FCS/PBS, with periodic gentle agitation.

Fixed and stained cells were pelleted by centrifugation (800 × g, 10 mins, room temperature), washed in PBS, re-pelleted and re-suspended in PBS for sorting by FACS.

The FACS sorting was conducted using FACSAria III (Becton Dickinson) instrument. The instrument was set up and QAd with SC&T beads prior to sort to achieve 99% sort purity with the beads. The parameters recorded were mCherry positivity and LTAg expression, with fluorescence detected in the channels 568 nm and 647 nm, using bandpass filters 660/20 and 610/20 for detecting lentiviral mCherry (568 nm) and LTAg (647 nm). After cells were identified via their morphological features (FSC and SSC), the population of mCherry positive and lowest 10% of LTAg expressing cells was sorted. The gates were set based on control untransfected cells.

These cells were pelleted by centrifugation (6,000 × g, 5 mins, 4°C), washed twice with PBS, further pelleted by centrifugation and stored at −80°C.

The reference library population was harvested by trypsinisation (10 mins, 37°C), pelleted by centrifugation (6,000 × g, 5 mins, 4°C), washed twice with PBS, further pelleted by centrifugation, and stored at −80°C, as per the sorted population.

Genomic DNA was extracted from both the sorted cells and the reference library population using the QIAamp DNA FFPE tissue kit following manufacturer’s instructions (Qiagen). The variable sgRNA sequences were amplified by nested PCR (Q5 High-Fidelity Polymerase, NEB), the first round of PCR amplifying all sgRNA insertions, and the second round of PCR amplified using primers with P5 (common) and P7 (specific) adaptors allowing subsequent Illumina sequencing (Table 1). The products of these PCR reactions were purified with Agencourt AMPure XP beads (Beckman Coulter) and pooled, allowing sequencing of the amplified products using a 50 bp single-end read on an Illumina HiSeq 2500 instrument.

**Table 1:**
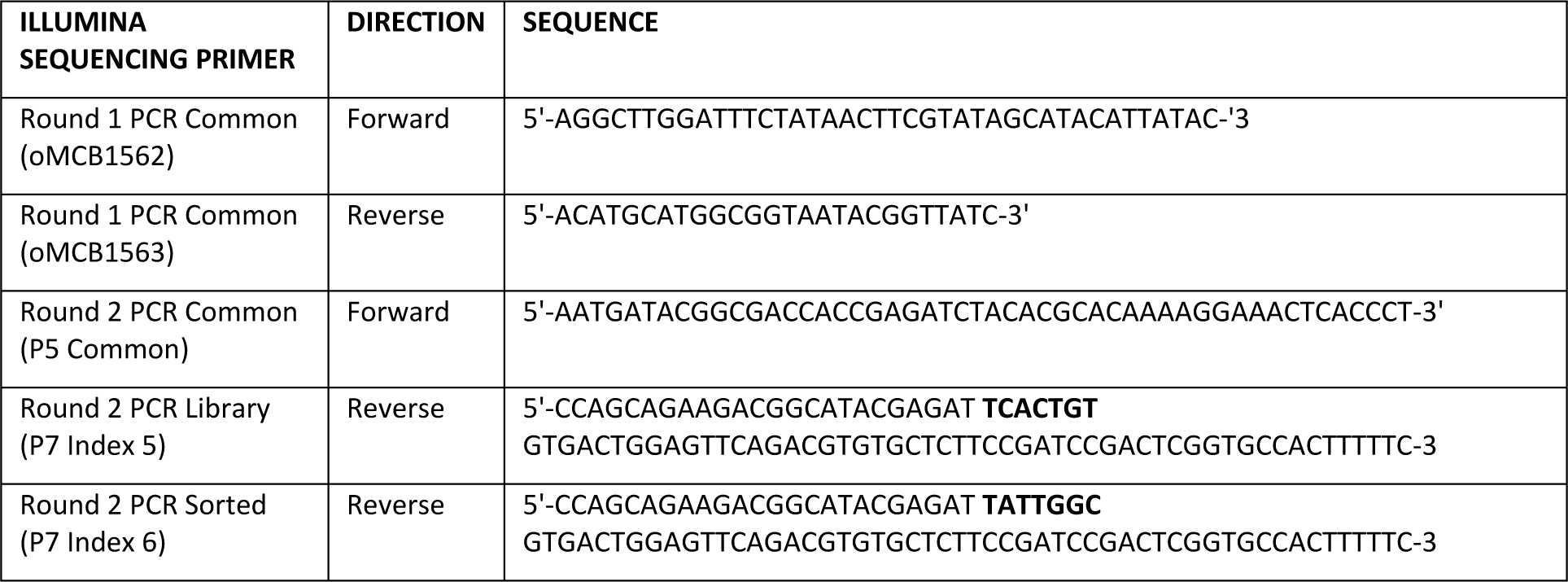
Illumina primers used in this study. Adapter sequences highlighted in bold.

### Illumina sequence read processing and analysis

Reads were 3’ trimmed with cutadapt v3.4 (Martin, 2011) to remove terminal Ns and adapters, keeping only trimmed reads between 15 and 26 nucleotides in length. Quality control was performed before and after trimming using FASTQC v0.11.9 (Andrews, 2010). Reads were then mapped to a FASTA file generated from the Bassik Human CRISPR Knockout Library (Addgene), clipped to remove one nucleotide at the 5’ end of each read (due to an excess of mismatches at this position). The length distribution of the trimmed reads was similar to that of the library (Figure S1Bi, ii, iii). Reads were mapped to the library with bowtie2 v2.4.2 (Langmead and Salzberg, 2012), with the default settings plus “--norc” to prevent mapping of reverse complemented reads. Mapping rates were consistently high, with 84-87% of the reads mapping to the library (Figure S1Biv). Read count tables were generated from the bam files using bowtie2 v2.4.2 idxstats (Langmead and Salzberg, 2012). Comparisons between conditions were performed with MAGeCK v0.5.9.4 (Li *et al*., 2014) using “control” normalisation (Figure S1Bv, vi). P-values per gene are taken from column pos|p-value in the MAGeCK gene level output table.

### Bioinformatic pathway analysis

The 200 most significantly enriched sgRNAs in the sorted population of this CRISPR screen were analysed using the DAVID Bioinformatics Resources (Version: DAVID 2021), performing a biological process and pathway analysis (Huang *et al*., 2007; Huang, Sherman and Lempicki, 2009), cut offs of an EASE score of 0.05, minimum 4 genes were implemented.

### Experimental Statistical analysis

Between experimental conditions significance was determined using a two-sample Student’s *t*-test.

---

**Figure S1:**
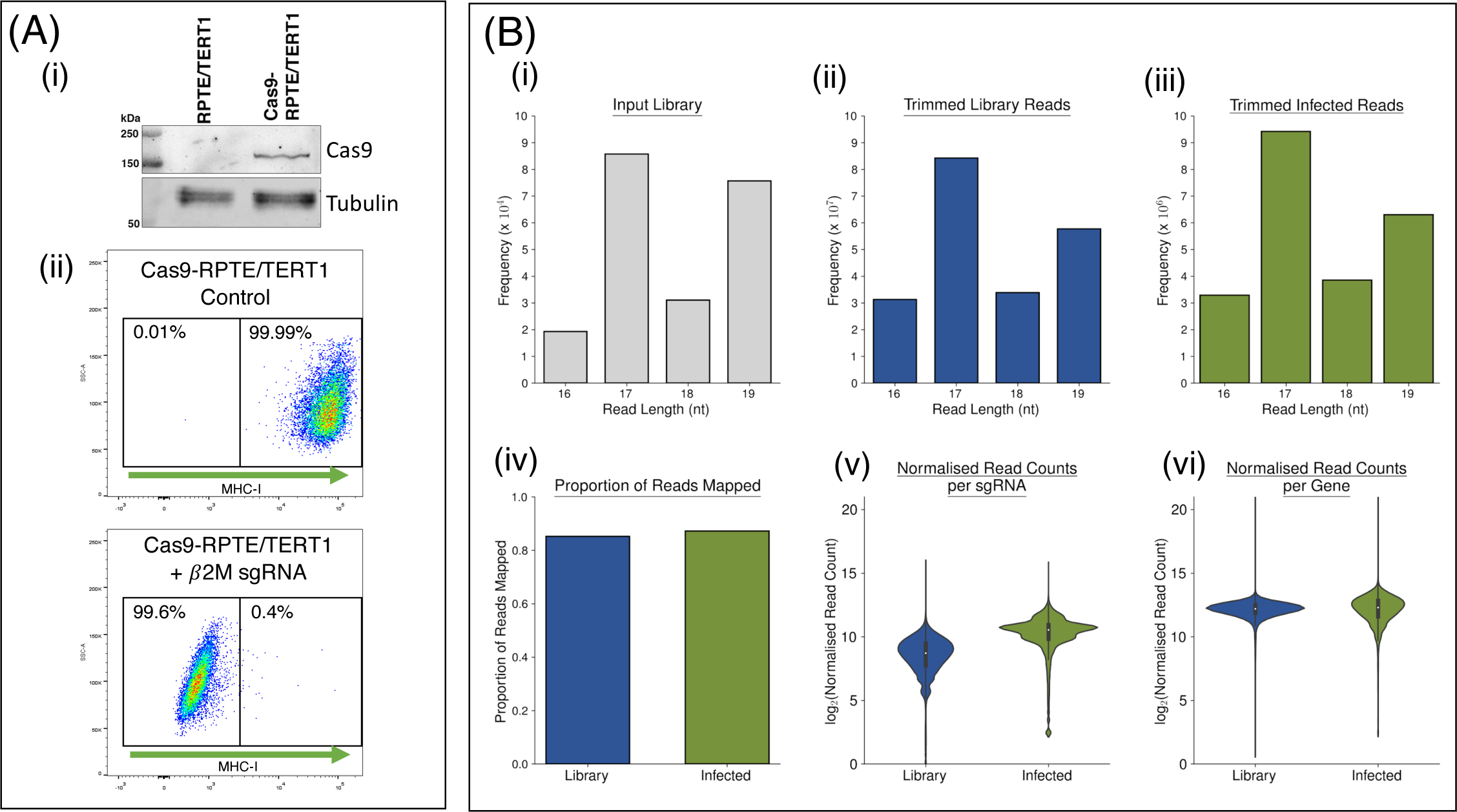
Cas9-RPTE/TERT1 Screen Validations. (A) *<u>Creation of Cas9-RPTE/TERT1 cell line</u>*: RPTE/TERT1 cells, a human renal cell line was lentivirus transduced to stably express Cas9, shown here by Western blot (i). Cas9 function was validated by further transducing β2M sgRNA, leading to reduction of MHC-I expression on the cell surface, shown here by FACS surface stain for MHC-I (ii). (B) *<u>Library Trimming and Quality Control</u>* (i-iii) Bar charts showing the distribution of read lengths; The Bassik Human CRISPR knockout library (Addgene), after clipping one nucleotide from the 5’ end of each read (i). The reads from the uninfected cells (ii). The reads from the infected cells, both after trimming terminal Ns and adapters (iii). Bar chart showing the proportion of reads successfully mapped to the library for uninfected (left) and infected (right) cells (iv). Violin plot showing the distribution of normalised read counts per sgRNA calculated with MAGeCK (Li et al., 2014) for uninfected (left) and infected (right) cells, excluding sgRNAs with counts of zero (v). Violin plot showing the distribution of normalised read counts summed per gene, calculated with MAGeCK (Li et al., 2014) for uninfected (left) and infected (right) cells, excluding genes with counts of zero (vi).

**Figure S2:**
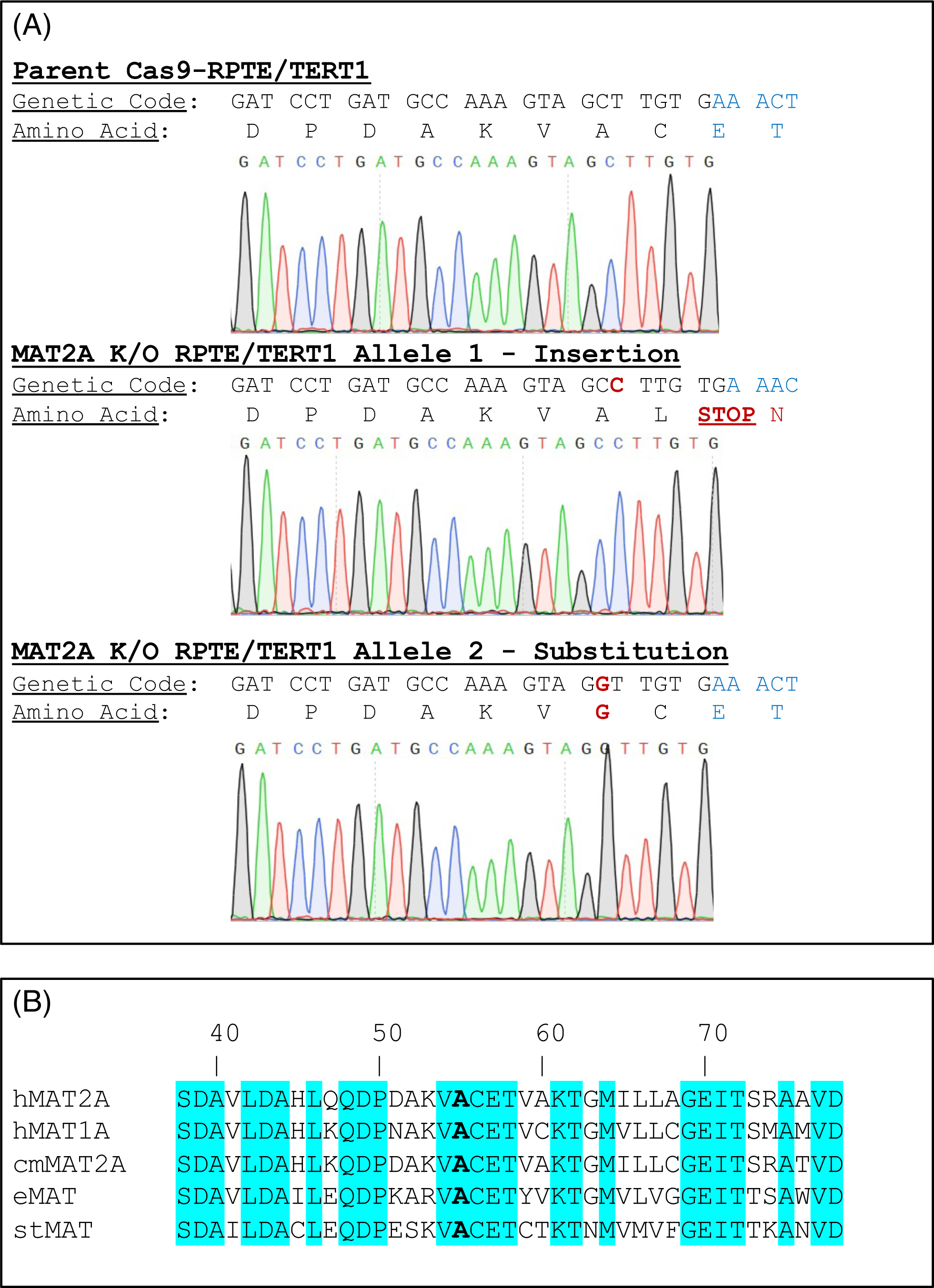
RPTE/TERT1 MAT2A K/O Indel Data. (A) Genetic code and amino acid code of MAT2A in parental Cas9-RPTE/TERT1 at Cas9 sgRNA cut site, compared with MAT2A K/O alleles. Points of difference in MAT2A K/O alleles shown in red. (B) Conservation of S-adenosylmethionine transferases across different domains. Residue numbering is for hMAT2A. Sequences given corresponding to homologs of hMAT2A (Human MAT2A Uniprot: P31153), hMAT1A (Human MAT1A Uniprot: Q00266), cmMAT2A (Callorhinchus milii [Ghost Shark] Uniprot: A0A4W3I8A4), eMAT1A (Escherichia coli MAT Uniprot: P0A817) and stMAT (Solanum tuberosum [Potato] Uniprot: Q38JH8). Conserved residues are highlighted in blue. Residue A55 (and residue equivalent in homologs), which is mutated to Glycine (G) on one allele of the MAT2A K/O cells used in this study, is highlighted in bold.

**Figure S3:**
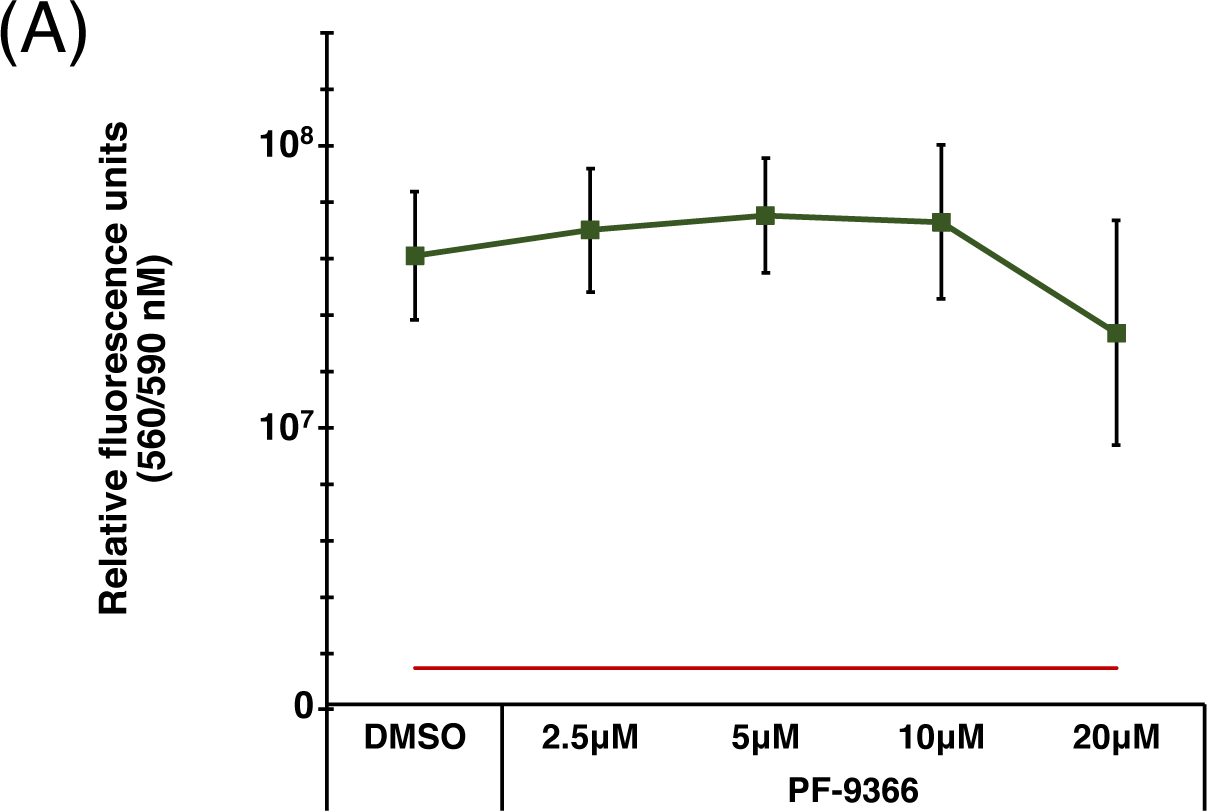
CellTiter Blue Metabolism Assay – PF-9366. *<u>Effect of PF-9366 on cellular metabolism</u>*: Cells were treated with PF-9366 at a range of concentrations for 48 hours and cell viability/metabolism was measured using CellTiter Blue assay. 60uM Roscovitine was used as a control, known to dramatically effect cell viability, giving a reading of 7.4E+06, shown as a red line for these purposes. n=3

**<u>Table S1: Complete Analysed HiSeq Data</u>**

Tab 1: All genes identified by HiSeq analysis, in alphabetical order

Tab 2: Filtered results, with those genes identified by less than 4 of the 10 sgRNAs used removed. Genes ranked by -LOG(P).

Tab 3: Filtered results, in alphabetical order for plotter

**<u>Table S2: DAVID Gene List</u>**

List of genes in each DAVID pathway identified.

## Notes

### Competing Interest Statement

The authors have declared no competing interest.

